# Evidence for an Intricate Relationship Between Express Visuomotor Responses, Postural Control and Rapid Step Initiation in the Lower Limbs

**DOI:** 10.1101/2022.10.21.513067

**Authors:** Lucas S. Billen, Brian D. Corneil, Vivian Weerdesteyn

## Abstract

Recent work has described express visuomotor responses (EVRs) on the upper limb. EVRs are directionally-tuned bursts of muscle activity that occur within 100 ms of visual stimulus appearance, facilitating rapid reaching. Rapid stepping responses are also important in daily life, and while there is evidence of EVR expression on lower limbs, it is unknown whether lower-limb EVRs are influenced by increased postural demands. Here, we investigate the interaction between stepping-related EVRs and anticipatory postural adjustments (APAs) that typically precede step initiation.

16 healthy young subjects rapidly stepped towards visual targets presented in front of the left or right foot. We recorded bilateral surface EMG of gluteus medius (GM), a muscle involved in both APAs and stepping, and bilateral ground reaction forces. Two conditions were introduced: an anterolateral or anteromedial stepping condition with reduced or increased postural demands, respectively. In the anterolateral stepping condition, EVRs were robustly and strongly present in stance-side GM, and ground reaction forces revealed strongly decreased expression of APAs. Larger EVRs preceded shorter RTs, consistent with EVRs facilitating step initiation. In contrast, in the anteromedial stepping condition, EVRs were largely absent, and ground reaction forces revealed the consistent expression of APAs. When occasionally present, EVRs in the anteromedial stepping condition preceded larger APAs and longer RTs. Thus, while EVRs in lower limbs can facilitate rapid stepping, their expression is normally suppressed when postural stability is low. Failing to appropriately suppress EVRs in such situations disrupts postural stability, necessitating larger compensatory APAs and leading to longer stepping RTs.

**Key Points:**

- Express visuomotor responses (EVRs) are directionally tuned bursts of muscle activity that aid the rapid initiation of a goal-directed movement. They are thought to be relayed to the motor periphery along a rapid subcortical pathway involving the superior colliculus.
- While EVRs have predominantly been studied in reaching, it is unclear whether EVRs extend to the lower extremities and if so, whether increasing the postural demands of a stepping task interfere with lower-limb EVR expression.
- We found that when postural demands were low, strong EVRs in the hip abductor muscle gluteus medius facilitated a rapid stepping response. Conversely, when postural demands were high, EVRs hindered a fast stepping response, as they necessitated larger, compensatory postural adjustments prior to step onset.
- These results help us better understand the interaction between ultra-rapid visuomotor transformations in the EVR network, the postural demands of a given stepping task, and subsequent step initiation.

## Introduction

In everyday life, extremely fast reaction times are frequently required to adequately respond to visual stimuli, for example when catching a ball that is suddenly thrown at us. Previous research has shown that our fast visuomotor system allows for reaction times of ∼120 ms in situations where mid-flight reaching adjustments were required (Day & Brown, 2001; Day & Lyon, 2000; Fautrelle, Ballay, et al., 2010; Fautrelle, Prablanc, et al., 2010; Soechting & Lacquaniti, 1983). A novel method that has been proposed to study the fast visuomotor system from a static starting position is through electromyographic (EMG) measurement of so-called *express visuomotor responses* (EVRs; formerly called ‘visual responses’ or ‘Stimulus-locked responses’; Corneil et al., 2004; Pruszynski et al., 2010). These are defined as short-latency bursts of muscle activity that occur in a time-locked window ≈100 ms after stimulus presentation and precede the larger volley of muscle activity associated with movement initiation. There are compelling similarities between the response properties of EVRs and the response properties of mid-flight corrections. For example, like on-line corrections (Day & Lyon, 2000), EVRs are directionally tuned toward the visual stimulus, even when the intended movement is in the opposite direction or temporarily suppressed (Atsma et al., 2018; Gu et al., 2016; Wood et al., 2015). Further, earlier and larger-magnitude EVRs are provoked by stimuli of high contrast or low spatial frequency (Kozak et al., 2019; Wood et al., 2015), similar to the response properties of on-line corrections (Kozak et al., 2019; Veerman et al., 2008). Evidence suggests that EVRs are relayed to the motor periphery along a subcortical tecto-reticulo-spinal pathway (Contemori et al., 2021b; Corneil & Munoz, 2014; Glover & Baker, 2019; Gu et al., 2017; Kozak et al., 2020; Pruszynski et al., 2010). This conclusion is based on the short latency of the EVR, its temporal separation from the larger wave of muscle recruitment associated with the voluntary movement, and the similarity in stimulus preferences with those seen in the visual responses in the intermediate superior colliculus (Chen et al., 2018; Marino et al., 2012; Rezvani & Corneil, 2008).

The vast majority of studies have focused on fast visuomotor responses of the upper limb. Fast stepping responses are arguably as important as fast reaching movements, for example when rapidly adjusting one’s stepping behavior while walking on uneven terrain or when intercepting a ball while playing soccer. Indeed, there is strong evidence that the fast visuomotor network can also recruit express responses in lower limb muscles, for instance during on-line pointing adjustments in an upright standing posture (Fautrelle, Prablanc, et al., 2010), or while making on-line stepping adjustments in response to an obstacle or a target shift (Nonnekes et al., 2010; Reynolds & Day, 2005). These on-line adjustments can be initiated substantially faster than voluntary stepping adjustments, with reaction times ranging from ≈105-150 ms (Marigold et al., 2007; Reynolds & Day, 2005; Weerdesteyn et al., 2004). Importantly, postural demands are substantially higher in stepping compared to reaching, because of the concurrent involvement of our legs in balancing the Centre of Mass (CoM). These findings suggest that a sudden visual stimulus triggers a rapid goal-directed adjustment of the stepping movement without integration of the postural demands, potentially destabilizing the body.

Here, we investigated during goal-directed stepping the interaction between the ultra-rapid EVR response in the lower extremities and postural control in the form of *anticipatory postural adjustments* (APAs). APAs typically precede step initiation from standstill and are closely tied to the size and direction of the ensuing step (Bancroft & Day, 2016; Inaba et al., 2020). To trigger EVRs, we used a recently developed emerging target paradigm that involves the sudden, but temporally predictable, appearance of a moving visual target below an occluder (see Figure 1). This paradigm has been shown to consistently evoke robust EVRs in upper limb muscles while making reaching movements from a static position (Contemori et al., 2021a; Kozak et al., 2020; Kozak & Corneil, 2021). It is thought that the use of implied motion behind a barrier, which has been shown to produce strong signals in motion-related areas in dorsal visual stream (Krekelberg et al., 2005), combined with the high certainty of the time of target appearance, results in a strong visual transient (Contemori et al., 2021a; Kozak et al., 2020). These properties have been shown to result in earlier, stronger, and more prevalent EVRs compared to other paradigms. In the current study, we modified this paradigm into a stepping task.

**Figure 1.**
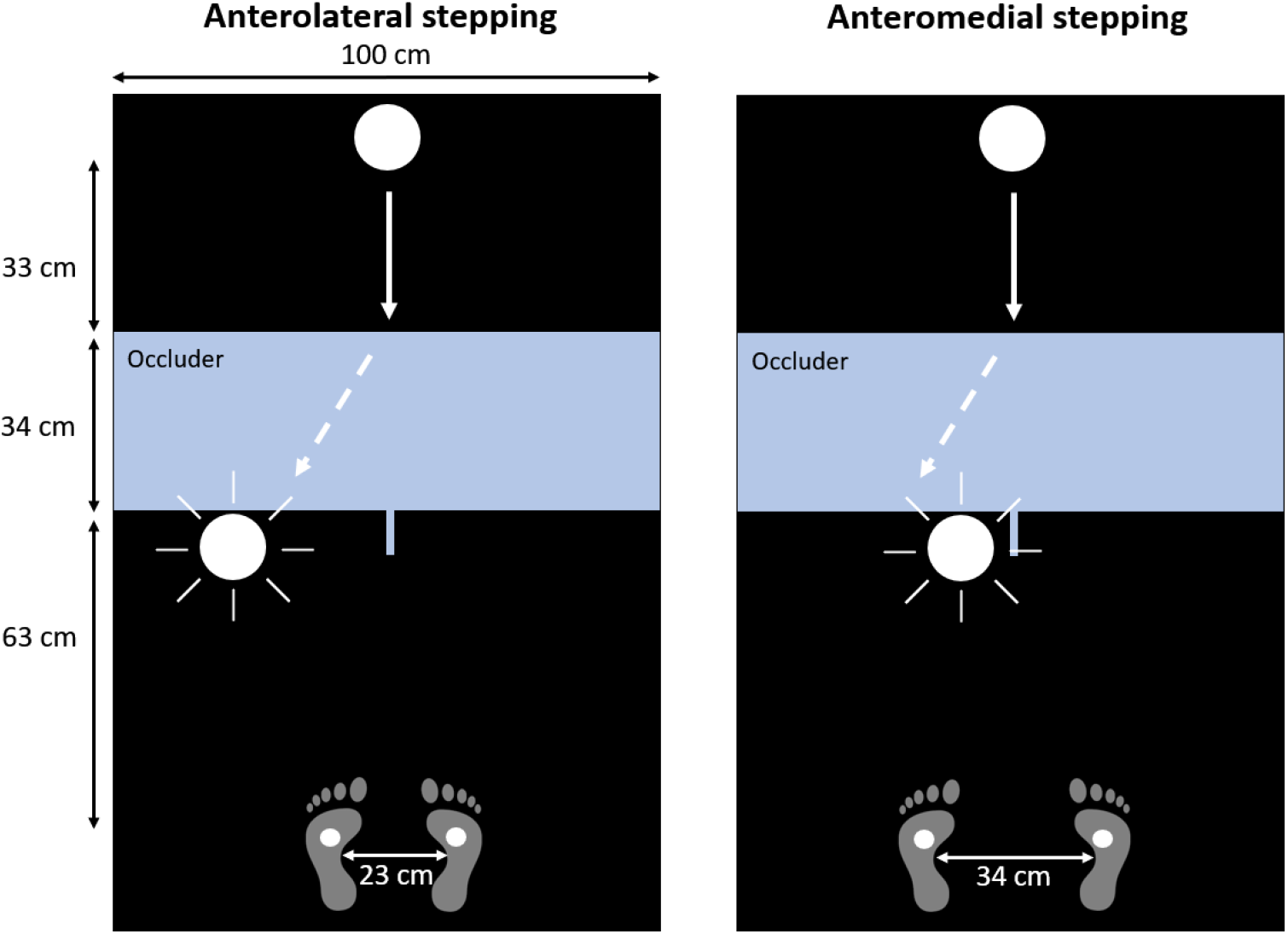
Experimental setup of the emerging target paradigm. The paradigm was projected on the floor in front of the participants. Participants placed their feet on two projected dots. The visual target moved down towards the participants, disappeared behind the occluder, and then, in this example, reappeared in front of left foot of the participant. Participant stepped onto the target upon reappearance, requiring either an anterolateral (left figure) or anteromedial (right figure) stepping response

We recorded EMG from two muscles, tibialis anterior (TA) and gluteus medius (GM), reasoning that they could be involved in both postural control and fast visuomotor responses. TA is commonly used to characterize APAs involved in step initiation. It activates bilaterally (yet more strongly in the stepping leg) during APAs to generate the initial forces that propel the CoM forward. GM is involved in both the APA and the ensuing stepping movement. APA-related GM activity is recruited on the *stepping* side to shift the CoM towards the stance side, followed by stepping-related activity in the *stance* leg to propel the CoM forward and towards the stepping side. Thus, GM in particular provides the unique opportunity to distinguish APA-related activity from stepping-related activity prior to step onset.

In order to investigate whether EVRs correspond to APA-related activity (i.e. ipsilateral GM activity in the stepping limb) or to stepping-related activity (i.e. contralateral GM activity in the stance limb), we introduced contrasting balance demands, across different blocks of trials. This was achieved by presenting the stepping targets in front of the participant either somewhat medially or somewhat laterally, and by varying initial stance width. Based on previous research demonstrating that stepping direction influences APA expression (Inaba et al., 2020), we expected that these task manipulations would necessitate strong APAs when subjects step anteromedially from a wider stance width (high postural demand during step execution), but would yield decreased APAs when subjects step anterolaterally from a narrow stance (low postural demand during step execution) (Bancroft & Day, 2016; Inaba et al., 2020). If, as in the upper limb, EVRs promote movement toward the target, then we expect EVRs to be generated in the *stance* side GM and possibly TA, as these muscles rapidly propel the body forward and toward the stepping side. Whereas such stance-leg EVRs may be beneficial for fast anterolateral stepping, strong stance-leg EVRs are expected to compromise postural stability in the anteromedial stepping condition, because they would counteract APAs initiated in stepping-leg GM, thereby compromising step initiation. Overall, we found that stance-leg EVRs were readily expressed when subjects stepped anterolaterally, but (when present) preceded larger APAs when subjects stepped anteromedially under greater postural demand.

## Materials and Methods

### Subjects

16 healthy young subjects (4 males, 12 females) participated in this study. Ages ranged from 19 to 28 years (*M* = 23.35, *SD* = 2.37). Only participants with a BMI under 25 kg/m^2^ were included in the study to minimize the coverage of muscles by adipose tissue, which could compromise the quality of surface EMG recordings, particularly of GM. None of the participants had any visual, neurological, or motor-related disorders that could influence their performance in the study. The study protocol was reviewed by the medical ethical committee (CMO Arnhem-Nijmegen, 2021-13269) and the study was conducted in accordance with the latest version of the Declaration of Helsinki. All participants provided written informed consent prior to participation and were free to withdraw from the study at any time.

### Experimental design

The experiment was performed using a Gait Real-time Analysis Interactive Lab (GRAIL, Motek Medical, The Netherlands). The experimental setup included an M-gait dual-belt treadmill with two embedded force plates (GRAIL, Motek Medical, The Netherlands) to measure ground reaction forces, a Wave Wireless electromyography system (Wave Wireless EMG, Cometa, Italy) to record muscle activity, and a projector (Optoma, UK) to project all visual stimuli. Participants stood on the stationary M-Gait with each foot placed on a separate force plate. The stepping task was projected on the treadmill in front of the participant (Figure 1). Previous research has demonstrated the importance of high-contrast stimuli in eliciting strong EVRs (Kozak & Corneil, 2021; Wood et al., 2015). We therefore covered the treadmill with a white vinyl mat and darkened the room. Additionally, all participants wore a cap that shielded their eyes from the light of the projector, which further increased the relative brightness of the projected stimuli. The stimuli had a luminance of 7 cd/m² against a black background of 0.23 cd/m² (contrast ratio: 30:1).

Each trial started with the appearance of a stationary visual target (solid white circle) 130 cm in front of the participant, presented on a black background. This distance was chosen based on previous research demonstrating that during walking, humans tend to fixate their eyes on the ground approximately two steps ahead (Patla & Vickers, 2003). After being visible for 1000 ms, the target started moving towards the participant at a constant speed of 0.44 m/s (with a retinal speed of ∼11.0°/s for an average sized participant of 169 cm). The target then disappeared behind an occluder (a light blue rectangle) and followed a straight track downwards to the base of the occluder. It remained invisible to the participants for a fixed interval of 750 ms. Because the target continued to move at a constant speed behind the occluder, participants could anticipate the timing of target reappearance; in the upper limb, such certainty of the time of target reappearance increases EVR prevalence, increases EVR magnitude, and decreases EVR latency (Contemori et al., 2021a). Once the invisible target had reached the base of the occluder, the target reappeared randomly in front of the left or right foot of the participant. Because this is the first study using this paradigm to study EVRs in the lower extremities, we varied target appearance to investigate its influence on EVR properties. The targets reappeared underneath the occluder as either (1) a target moving towards the lateral edges of the treadmill at a constant speed of 0.23 m/s, or (2) a flashed target (one single flash with a duration of 48 ms (i.e. three frames)). We realized in retrospect that moving targets introduce a confound between reaction time and stepping length (since moving targets move away from the subject). Moreover, we aim to use this paradigm for comparing EVRs between neurological patients and healthy controls. The interpretation of between-group comparisons is more straight-forward when the behavioral response (i.e. step length) is kept constant, therefore In the present article, only results for the flashed condition will be reported. We found only minor differences in EVR expression between the flashed and the moving condition. These do not influence our interpretation of the results (see Supplementary Materials).

Participants were instructed to divide their weight equally between both legs prior to step onset and to avoid shifting the CoM forward in anticipation of the reappearing target. This was visually checked during the experiment based on real-time force plate data. Participants were instructed to perform a full stepping movement upon reappearance of the target, using the leg on the side of target appearance (i.e. step with the left leg when the target appeared on the left side and vice versa for the right leg). After having stepped onto the target with the stepping leg, the stance leg had to be placed next to the stepping leg to complete the stepping movement. It was emphasized to the participants that speed was the most important parameter in the present study and that the step had to be initiated as rapidly as possible. After having completed the trial, the participant returned to the starting position and the subsequent trial was initiated.

Trials were started manually via the D-flow software (Motekforce Link, The Netherlands) by the experimenter. To account for small variable delays in target presentation, a photodiode (TSL250R-LF, TAOS, USA) was used to measure the exact moment of target appearance. This was achieved by placing the diode over a secondary peripheral target presented at the same time as the actual stepping target. This secondary target was presented outside of the participant’s field of view. All reported measures (i.e. EMG and force plate measures) were aligned to the moment of stimulus presentation detected by the photodiode.

In order to investigate the interaction between postural control in the form of APAs and EVRs, targets were presented in front of the stepping foot, either anterolaterally or anteromedially. The primary stepping direction for both the anterolateral and anteromedial condition is forward, as the forward stepping length is 63 cm. In the anterolateral target condition, participants started at a narrow stance width (feet 23 cm apart) and stepped forward and outward towards an anterolateral target presented 29 cm from the middle line of the treadmill (retinal eccentricity ∼10°; stepping eccentricity ∼11.6° outward relative to straight ahead from stepping foot). In the anteromedial target condition, participants started at a wide stance width (feet 34 cm apart) and stepped forward and inward towards an anteromedial target presented 9 cm from the middle line (retinal eccentricity of ∼3°; stepping eccentricity ∼8.1° inward relative to straight ahead from stepping foot). The variation in stance width between the anteromedial and anterolateral stepping condition was used to increase the relative contrast with regard to stepping direction and with regard to balance control: stepping medially from a wide stance width increases the balance demands, and consequently the need to make an APA, while stepping laterally from a narrow stance width on anterolateral targets has the opposite effect. In this context, it is important to highlight that the manipulation of stance width and target location entails the simultaneous manipulation of two distinct variables. The main objective in doing so was to maximize the contrast in balance demands between the two conditions.

Participants completed 4 blocks of 150 trials (600 in total). Each block consisted of either only anterolateral targets or anteromedial targets and the order of the blocks was counterbalanced.

Participants were informed about the condition before each block. Target appearance (moving/flashed) and target side (left/right) were randomized on each trial. Participants started with a few practice trials to become familiar with the task. The initial stance position was indicated by the projection of small circles at the desired foot location.

### Data collection

We recorded muscle activity from bilateral TA and GM using Ag/AgCl surface electrodes placed approximately 2 cm apart and longitudinally on the belly of the muscle (Wave Wireless EMG, Cometa, Italy). Skin preparation and electrode placement were performed in accordance with the SENIAM guidelines (Hermens & Merletti, 1999). The quality of the signal was checked before starting the recording session. EMG and force plate data were sampled at 2000 Hz.

### Data processing and analysis

Incorrect trials were excluded from the analysis. Incorrect trials were defined as trials in which participants stepped towards the wrong direction or initiated stepping movement with the contralateral foot. Data analysis was performed using custom-written MATLAB scripts (version 2019a).

#### Reaction time

Stepping RT was defined as the time from visual target appearance, as measured by the photodiode, to the foot-off moment of the stepping foot. In line with previous research, foot-off was defined as the first sample at which the vertical ground reaction force component (Fz) was lower than one percent of the participants body weight (Rajachandrakumar et al., 2017).

#### EVR presence and latency

The raw EMG signals were first band-pass filtered between 20 and 450 Hz and subsequently rectified and low-passed filtered at 150 Hz. Second-order Butterworth filters were used. To determine the presence and latency of EVRs, we used a time-series receiver-operating characteristic (ROC) analysis, as described previously (Gu et al., 2016; Kozak et al., 2020). EMG data were grouped based on target location (lateral vs medial), and target side (left vs right). EMG activity of GM and TA was then compared between leftward and rightward steps within any condition. For every sample between 100 ms before and 500 ms after visual stimulus appearance, an ROC analysis was performed and the area under the ROC curve (AUC) was calculated. This metric indicates the probability that an ideal observer can discriminate between the sides of stimulus location based solely on EMG activity. The AUC values range between 0 and 1, where a value of 0.5 indicates chance discrimination and values of 1 or 0 indicate perfectly correct or incorrect discrimination, respectively. Differences in muscle recruitment on leftward vs rightward steps is thus crucial to ensure robust EVR detection. In line with previous research, we set the discrimination threshold to 0.6 (Gu et al., 2016). The time of earliest discrimination was defined as the time at which the AUC surpassed the discrimination threshold and remained above the threshold for 16 out of 20 consecutive samples within the pre-defined EVR epoch of 100-140 ms after stimulus presentation. Compared to reaching studies, which typically used EVR windows of 80-120 ms (e.g. (Gu et al., 2016; Kozak et al., 2020)), the epoch used in the current study was adjusted based on the use of lower contrast stimuli (Kozak & Corneil, 2021), the conduction velocity of reticulospinal neurons and the additional distance to the lower extremities (Buford, 2009).

#### Response magnitude in EVR window

The response magnitude in the EVR window was calculated for each condition within each participant, regardless of whether an EVR was detected. On a single trial basis, the average EMG activity during the EVR epoch (100-140 ms) was calculated and normalized against the median peak EMG activity (in the interval from 140 ms to foot-off) within the moving target condition during anterolateral stepping of the respective subject. EMG magnitudes of all trials were then averaged per condition.

### Statistical analysis

Statistical analyses were performed using IBM SPSS statistics software (version 27). The level of significance was set to *p* < .05 for all analyses. We used paired t-tests to investigate differences in EVR discrimination times between left and right muscles. Further t-tests were performed to study whether EVR magnitude and subsequent stepping RT differed between anterolateral and anteromedial stepping. To further investigate the relationship between EVR magnitude and subsequent stepping RT, we determined Spearman’s rank correlation coefficients on the single trial data of all participants.

APA magnitude was defined as the maximum vertical ground reaction force component (Fz) underneath the stepping leg in the interval from 140 ms after target appearance (i.e., the end of the EVR window) and foot off, normalized to percent total body weight. To establish APA onset, a sliding one-sample t-test was performed to test for any given time-point if the mean force plate data (grouped by condition (lateral/medial) and stepping side (stepping/stance side)) significantly deviates from the respective baseline value that was chosen to be at 100 ms after stimulus presentation. The first of at least 10 consecutive statistically significant samples was defined as APA onset.

In order to further characterize the relationship between EVR presence and subsequent stepping RT, trials were split based on median RT, thereby creating a “fast RT” and “slow RT” subset. Subsequently, time-series ROC analyses were performed on each subset separately to determine EVR presence within the two subsets. The same criteria for EVR presence as described above were applied.

## Results

### Incorrect trials

The overall rate of incorrect trials was low; 1.06% of all steps involved errors in the anteromedial condition and 0.43% of all steps involved errors in the anterolateral condition. Error rates differed greatly between subjects. Two subjects made only a single error out of 600 trials (error rate: 0.16%). In contrast, the most error prone subject made 58 errors in total (error rate: 9.7%).

### Visual Inspection of EMG Data Indicates the Presence of EVRs

Figure 2 shows muscle recordings of an exemplar participant aligned to visual stimulus onset. The EMG signal of this participant showed one of the highest signal-to-noise ratios and exemplifies key features of the recruitment patterns of interest (see supplementary materials for data from other subjects). The first column shows the mean EMG activity of left and right GM (top rows) or TA (bottom rows) on anterolateral and anteromedial steps respectively. In addition, the time-series ROC curve is plotted above the EMG activity; the times at which this curve goes above/below the 0.6/0.4 thresholds indicates consistent differences in muscle recruitment when the associated leg acts as the stance or step limb.

**Figure 2.**
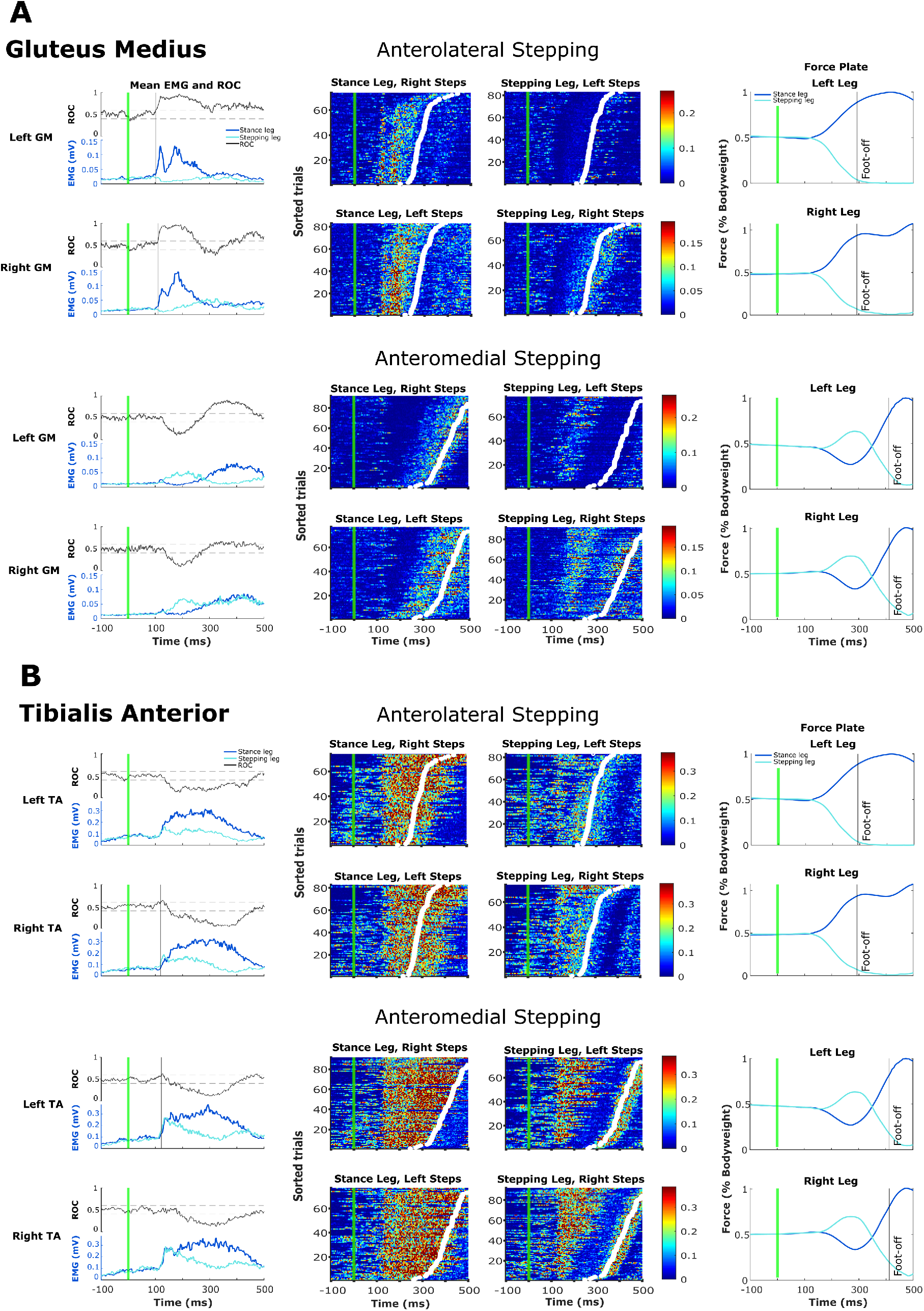
GM (A) and TA (B) muscle activity, time-series ROC analysis and force plate data of an exemplar participant. Data is separated based on muscle side (left/right) and stepping condition (anterolateral/anteromedial). Each condition is presented on a separate row. All data are aligned to visual stimulus onset (green line). **Column 1:** shows mean EMG activity for the stance-side (dark blue) and the stepping side (light blue) of the respective muscle. The time-series ROC curve is shown in gray. Discrimination times within the SLR epoch (100-140 ms) are indicated by the black vertical line. **Columns 2 and 3:** Trial-by-trial EMG activity of the stance side (column 2) and the stepping side (column 3). Intensity of color conveys the magnitude of EMG activity Each row represents a different trial. Trials are sorted by RT (white dots). **Column 4:** Mean vertical force (F_z_) exerted by the stance (dark blue) and stepping leg (light blue).

The first two rows of Figure 2A, which demonstrate GM activation on anterolateral steps, indicate that GM is primarily recruited on the stance limb side (e.g. left GM is active when stepping rightward). There are two distinct bursts of activity; the first peaks at around 110 ms and the second one peaks at around 190 ms after stimulus onset. These activation patterns are also visible in the trial-by-trial representations of recruitment, for which the associated muscle is on the stance (column 2) or stepping (column 3) limb. As can be inferred from the trial-by-trial activity in column 2, there is a clear burst of activity ∼110 ms after target presentation in almost every trial on the stance limb. This initial burst of activity is followed at a variable interval by a second longer-lasting burst that persists until shortly before foot-off (white dots). In contrast, column 3 shows that GM activity on the stepping limb is suppressed at ∼120 ms after target presentation and essentially remains silent throughout the trial. The initial burst of activity observed on the stance side is the EVR, as it is time-independent of the subsequent stepping reaction time. The ROC curve (column 1) supports this notion as the 0.6 threshold is crossed at 102 ms and 110 ms for left and right GM, respectively.

The kinetic consequences of these recruitment patterns are shown in the force plate data in column 4. From this, it is clear that ground reaction forces increase and decrease on the stance and stepping side, respectively, soon after the EVR (∼140 ms after target appearance). This presentation is important because it indicates that APAs were not expressed in the anterolateral stepping condition, given the absence of an initial increase in vertical forces on the stepping side.

The activation patterns on GM and the force plate data are distinctly different in the anteromedial stepping condition. As expected, APAs were clearly expressed in this condition, as shown by the initial increase in vertical forces on the stepping side at ∼150 ms, which potentially induced a CoM shift towards the stance side before foot lift off (see column 4).

Further, the underlying pattern of GM activation in the anteromedial condition exhibited a tri-phasic recruitment of stance- and stepping-leg GM. Mean EMG traces show that GM on the stance limb side was very briefly activated ∼100 ms after stimulus presentation, but was then immediately silenced. Subsequently, stepping-leg GM (e.g. left GM when stepping leftward) became active at approximately 130-300 ms after stimulus presentation, which is the timing and patterning expected of an APA. After 250 ms, GM on the stance limb became highly active and remained so through foot-off.

Figure 2B shows TA activity of the same exemplar subject. Overall, and in contrast to what was observed on GM, the initial activation pattern for both the anterolateral and the anteromedial stepping conditions looks rather similar. TA is symmetrically activated on both the stance and the stepping side starting at around 120 ms after stimulus presentation. As can be inferred from the ROC curve (column 1), a discrimination time within the pre-defined EVR window is absent in TA, due to this symmetrical activation. On the stance side, TA activation is maintained through foot-off, whereas on the stepping side, this initial activation is subsequently inhibited until shortly before foot-off. While at first glance it seems that TA on the stepping limb shows EVR-like activity, given that the initial recruitment is more time-locked to stimulus rather than movement onset, note that TA on the stance limb is being recruited at the same time. Thus, unlike GM in the anterolateral stepping condition, the recruitment of TA lacks the lateralization of recruitment to one limb or another that is a defining characteristic of an EVR.

### Robust and postural-dependent expression of EVRs on GM but not TA

As is shown in Table 1, EVRs were robustly detected on stance-side GM in the anterolateral condition for both left (16/16 participants) and right GM (15/16), whereas only a small number of participants exhibited EVRs in the anteromedial condition (1/16 for left GM, 3/16 for right GM). Discrimination times (see Table 1) did not differ significantly between left and right GM during anterolateral stepping (p > .1). Discrimination times on anteromedial steps were on average later compared to anterolateral steps (Table 1). Statistical analysis was, however, not possible due to the low number of participants who expressed EVRs in the anteromedial condition. Because the comparisons of both discrimination times and EMG magnitudes in the EVR window between right and left GM yielded similar results, and for reasons of conciseness, we chose to only report results for left GM for the remainder of the present paper.

**Table 1.** Discrimination times for GM and TA across participants in anterolateral vs anteromedial stepping (in ms). Empty cells indicate no discrimination time in that condition.

| Participant | Left GM |  | Right GM |  | Left TA |  | Right TA |  |
| --- | --- | --- | --- | --- | --- | --- | --- | --- |
|  | Lateral | Medial | Lateral | Medial | Lateral | Medial | Lateral | Medial |
| 1 | 107 | - | 100 | - | 130 | - | 119 | 134 |
| 2 | 101 | - | 102 | - | 127 | 138 | - | - |
| 3 | 105 | - | 107 | - | - | 117 | - | - |
| 4 | 102 | - | 110 | - | 136 | - | - | - |
| 5 | 103 | - | 111 | - | - | - | - | - |
| 6 | 106 | - | 100 | 118 | - | - | - | - |
| 7 | 106 | - | - | - | 133 | 134 | 130 | - |
| 8 | 115 | - | 112 | - | 130 | - | 140 | - |
| 9 | 103 | - | 102 | - | - | - | 139 | 121 |
| 10 | 115 | - | 113 | - | - | - | - | - |
| 11 | 101 | - | 106 | 107 | - | - | - | - |
| 12 | 122 | - | 113 | - | 129 | 122 | - | - |
| 13 | 110 | - | 105 | 111 | 121 | - | - | - |
| 14 | 109 | 113 | 103 | - | - | - | 136 | 104 |
| 15 | 106 | - | 103 | - | - | - | 138 | - |
| 16 | 119 | - | 107 | - | - | - | 131 | - |
| Mean | 108 | 113 | 106 | 112 | 129 | 127 | 135 | 120 |
| SD | 7 | - | 5 | 6 | 5 | 10 | 5 | 15 |

Although an increase in TA activity was commonly observed in the EVR time window, initial TA activation was often symmetrical, thus not yielding a discrimination time in the EVR window. As a result, we detected EVRs in TA much less consistently compared to GM. The discrimination times for TA were also considerably longer than for GM, and we suspect that they result from the lateralization of muscle activity that blended into the later part of the EVR time window. TA is therefore not a suitable candidate for studying EVRs in the context of this study, hence remaining analyses will focus on GM. Note that this does not exclude possible EVR detection in other tasks or different initial postures that might lead to a lateralization of TA activity.

### Magnitude of activity in the EVR window is higher in anterolateral stepping compared to anteromedial stepping

Consistent with the observations from visual inspection of the EMG data (Figure 2) and the absence of detectable EVRs in the anteromedial condition, the magnitude of EMG recruitment in the EVR interval was significantly higher in the anterolateral condition (*M* = 0.12 AU, *SD* = 0.05) compared to the anteromedial condition (*M* = 0.05 AU, *SD* = 0.02; *t*(15) = 6.12, *p* < .001, Hedges’ *g* = 1.50).

### Stronger EVRs precede shorter stepping reaction times

We investigated whether the contrasting postural demands of anterolateral and anteromedial stepping had an impact on stepping reaction times. Indeed, stepping reaction times were significantly shorter in the anterolateral condition (*M* = 314 ms, *SD* = 61 ms) compared to the anteromedial condition (*M* = 443 ms, *SD* = 54 ms; *t*(15) = 26.23, *p* < .001, *Hedges’ g* = 4.58; see Figure 3B). As previous work demonstrated that stronger EVRs in a reaching paradigm precede short reach RTs, we evaluated whether this also applied to our stepping paradigm. Indeed, in anterolateral stepping there was a strong negative correlation between EVR magnitude and subsequent stepping RT (*r* = - .582, *p* < .01), indicating that stronger EVRs preceded faster stepping RTs. This correlation was absent in the anteromedial stepping condition (*r* = -.040).

**Figure 3.**
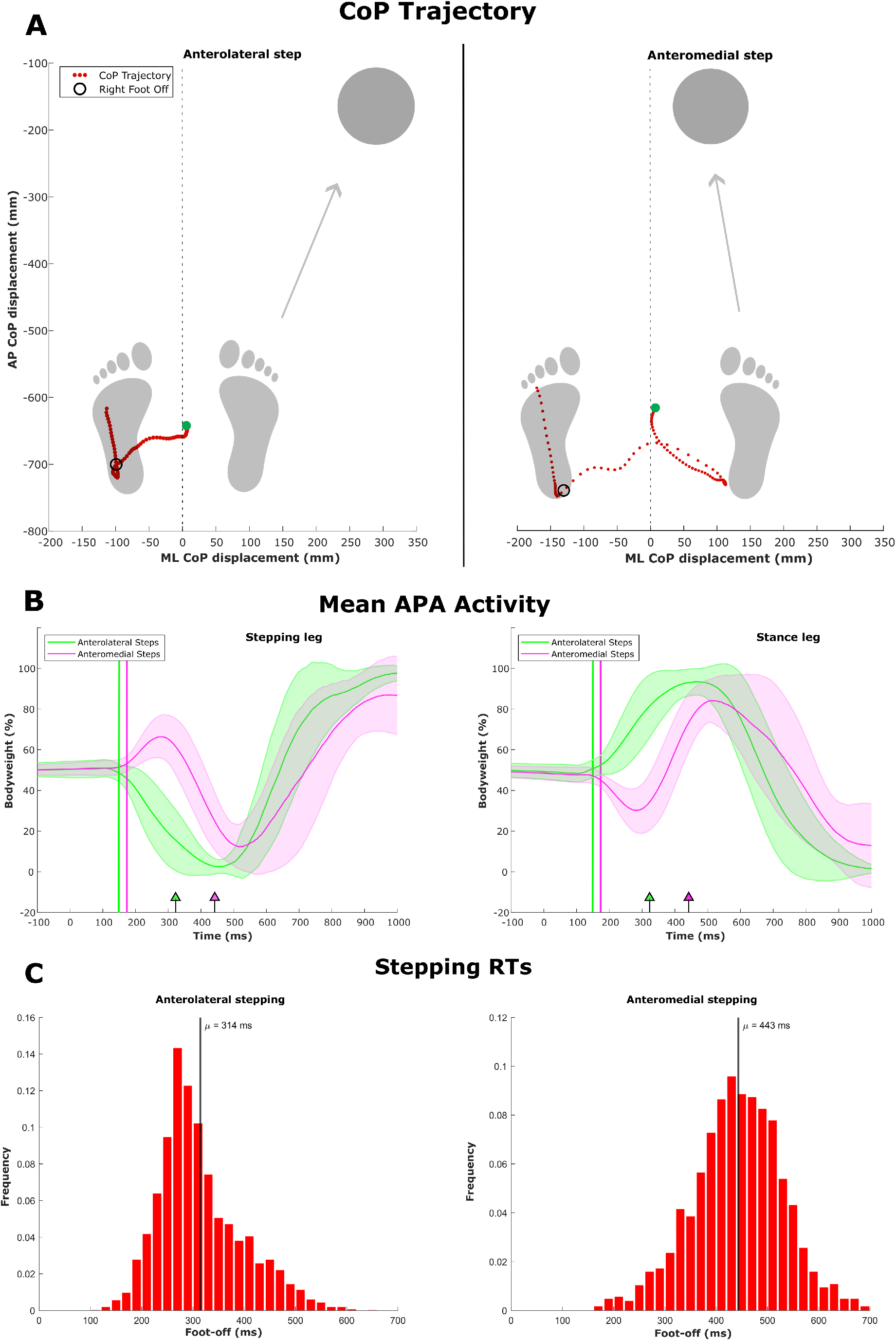
**A:** CoP trajectory of two representative trials in the anterolateral (left side) and anteromedial (right side) stepping condition (5ms spacing between dots). The green dot indicates the moment of target onset. The trajectory ends at the moment that the stepping foot (the right foot in these examples) lands on the target. Figure is drawn to scale regarding feet position and target location. **B:** Mean APA activity based on force plate data in anterolateral and anteromedial stepping underneath the stepping leg (left) and the stance leg (right). Shaded error bars indicate +/- 1 SD dispersion. Vertical lines indicate when the force plate data starts to significantly deviate from baseline, indicating APA onset. Arrows indicate mean stepping reaction times. **C**: Histogram of the trial-by-trial stepping reaction times of all subjects in the anterolateral (left) and anteromedial (right) stepping condition.

### Consistent APA expression in anteromedial stepping, but not in anterolateral stepping

Based on previous studies investigating the relationship between APAs and stepping eccentricity (Bancroft & Day, 2016; Inaba et al., 2020), we hypothesized that APAs would be expressed in the anterolateral stepping condition, albeit with a strongly decreased magnitude compared to anteromedial stepping. However, the rapid average stepping reaction times of 314 ms in the anterolateral stepping condition leave very little time to perform a complete APA. In addition, the representative subject described above suggests that APAs may not be expressed altogether in the anterolateral stepping condition. We therefore investigated the expression of APAs in the anterolateral and anteromedial condition.

Figure 3A illustrates the trajectory of the Center of Pressure of two representative trials from the aforementioned subject. The left subfigure displays a trial performed in the anterolateral stepping condition. The instantaneous shift of the CoP trajectory towards the stance side (left side) indicates that the stepping leg is immediately unloaded upon target appearance, which suggests the lack of anticipatory postural adjustment (APA) expression. The right subfigure displays a trial from the anteromedial stepping condition. Notably, in contrast to the anterolateral stepping condition, the Center of Pressure initially shifts towards the stepping side (right side). This CoP excursion implies that the stepping leg is actively generating forces to shift the center of mass towards the stance side, thereby subsequently allowing the stepping leg to be lifted off the ground to complete the step.

Figure 3B shows the mean vertical forces across all participants in the anterolateral and anteromedial stepping conditions when stepping towards the right side. As can be inferred from the magenta lines (i.e. anteromedial steps) in Figure 3B, and consistent with the force plate data and CoP trajectories shown for the representative subject, the small peak in vertical force underneath the stepping leg and the dip in vertical force underneath the stance leg indicate the typical expression of APAs. We found that the vertical force underneath the stepping leg started to exceed the baseline force at 172 ms after stimulus presentation. In the anterolateral condition (green lines in Fig 3), vertical forces immediately started to decrease underneath the stepping leg and increase underneath the stance leg, indicating that, strikingly, APAs were not only decreased in magnitude, but generally not expressed in this condition. The mean force started to significantly deviate from baseline at 150 ms in both stance and stepping leg, consistent with a shift of body weight towards the stance leg and unloading of body weight over the stepping leg.

### Relationship between EVRs, APAs and subsequent stepping RT

#### EVRs precede slow-RT steps in the anteromedial condition

As shown above, APAs are clearly expressed in the anteromedial condition. Interestingly, as shown in Fig. 2, the timing of EVRs on the stance side in the anterolateral condition preceded the timing of activity associated with APAs on the stepping side in the anteromedial condition. These findings led us to wonder if stance-side EVRs, if they were they produced in the anteromedial stepping condition, would influence APAs and overall subsequent stepping behavior. Although EVRs were indeed mostly absent in the anteromedial condition when performing the time-series ROC analysis for the whole set of trials, we frequently observed EVR-like activity on the slower half of trials upon visually inspecting the single trial data of all subjects. We therefore aimed to systematically investigate if EVRs indeed precede slower stepping RTs by performing separate time-series ROC analyses on the fast and slow RT subsets of trials, respectively. As shown in Table 2, EVRs were detected in no participants on the fast subset of trials (2^nd^ column in Table 2). In contrast, 9/16 participants exhibited EVRs on the slow half of trials (3^rd^ column in Table 2). Thus, EVRs regularly preceded steps with a subsequent slow stepping reaction time, suggesting that EVR expression in this condition potentially compromised the subsequent stepping behavior.

**Table 2.** Discrimination times indicating EVR presence across participants in the anteromedial stepping condition for the fast and the slow half of trials (in ms). Empty cells indicate no discrimination time in that condition. The last two columns show correlation coefficients and level of significance between trial-by-trial EVR magnitudes and subsequent APA magnitudes (based on force plate data) on the slow half of trials in the anteromedial stepping condition. * Significant at the .05 level. ** Significant at the .001 level.

| Participant | Anteromedial Stepping |  | Correlation EVR x APA |  |
| --- | --- | --- | --- | --- |
|  | Fast half | Slow half | *r* | *p* |
| 1 | - | - | .112 | .534 |
| 2 | - | 100 | .320 | .061 |
| 3 | - | 108 | .474* | .002 |
| 4 | - | - | .286 | .054 |
| 5 | - | 116 | .585** | < .001 |
| 6 | - | - | .408* | .018 |
| 7 | - | 113 | .386* | .020 |
| 8 | - | 113 | .038 | .823 |
| 9 | - | 107 | .318 | .062 |
| 10 | - | - | .587** | < .001 |
| 11 | - | - | .447* | .0003 |
| 12 | - | 102 | .360* | .024 |
| 13 | - | - | .250 | .120 |
| 14 | - | 113 | .229 | .185 |
| 15 | - | 114 | .114 | .522 |
| 16 | - | - | .463* | .008 |

If, indeed, stance-side EVRs interfered with step initiation in situations with high postural demands, then the disruptive consequences of stance-side EVR expression might have to be compensated for by larger stepping-side APAs. We therefore investigated the relationship between trial-by-trial EVR magnitude on the slow half of trials in the anteromedial stepping condition and subsequent APA magnitude (measured by the maximum vertical force underneath the stepping leg). Indeed, we overall observed positive correlations between EVR and APA magnitude, which were significant in eight participants, with stronger EVRs generally being followed by larger APAs.

## Discussion

In this study, we investigated the relationship between postural control and express visuomotor responses in tibialis anterior and gluteus medius using an emerging target paradigm. Participants performed fast stepping movements towards projected targets on the floor. These targets were presented anterolaterally or anteromedially relative to either a narrow or wide stance, respectively. In line with our hypotheses, we found that the emerging target paradigm could robustly evoke EVRs in GM. The EVRs corresponded to the stepping movement, meaning stance-side GM was active within the EVR window, while stepping-side GM activity was virtually absent. In the anterolateral stepping condition, this pattern of EVR expression facilitated the rapid execution of the step to propel the body forward and laterally, as stronger EVRs were followed by shorter stepping reaction times. In contrast, EVRs were largely absent in the anteromedial condition, but when present, they preceded larger APAs and longer step initiation times. Together, our findings point towards an intricate relationship between EVRs, APAs, and step initiation. EVRs precede APAs, and can potentially be suppressed in a posturally-dependent fashion. Whenever this suppression temporarily lapsed in a posturally-unstable condition, the EVR on the stance leg perturbed balance, necessitating a stronger, longer-lasting, and presumably compensatory APA on the stepping leg, which ultimately lead to longer reaction times.

### EVR characteristics are in line with previous studies

#### EVR Prevalence

We consistently observed early target-directed activity in the GM abductor muscle across all participants starting at a latency of ≈100ms after stimulus presentation. The short latency and time-locked nature of the observed GM activity are in line with the definition of an EVR as proposed by previous reaching studies (e.g. Contemori et al., 2021a; Glover & Baker, 2019; Gu et al., 2017; Kozak et al., 2019; Pruszynski et al., 2010). Thus, we here demonstrated that the emerging target paradigm that so efficiently introduced the cognitive factors necessary for EVR expression in the upper limbs (Kozak et al., 2020), can be adapted to a stepping paradigm leading to equally robust EVR expression in the lower limbs.

#### EVR Latencies

The average EVR latency was 107ms (ranging from 100-122 ms across subjects) and was similar across all participants and conditions, regardless of stepping reaction time, which underlines the time-locked nature of the EVRs. Compared to EVRs on upper limb muscles, the reported latencies are consistent with neural signals needing more time to travel to lower limb. Indeed, the latencies found in this study fit with the earliest EVR discrimination time in the pectoralis major muscle of 80 ms (Gu et al., 2016; Kozak et al., 2020; Contemori et al., 2021a; Kozak & Corneil, 2021) plus an additional average neural signal traveling time to the lower extremity of 20-25 ms via the reticulospinal tract (Buford, 2009). EVR latencies also depend on stimulus contrast, with high-contrast stimuli promoting earlier EVRs (Kozak & Corneil, 2021; Wood et al., 2015). This use of lower contrast stimuli in the current study compared to previous reaching studies may also have led to the longer EVR latencies reported here. In general, the EVRs in our study preceded the burst of muscle activity associated with voluntary movement, except for some cases in which the EVR fused with the voluntary stepping activity, especially on trials with shorter RT. Importantly, as observed during anteromedial stepping, EVR latencies also preceded the onset of APA-related muscle activity on the stepping side by ∼60 ms.

#### EVR Magnitudes

Another defining feature of the EVR is how its magnitude is inversely proportional to movement reaction time (Corneil et al., 2004; Pruszynski et al., 2010), consistent with the EVR imparting behaviorally-relevant forces (Gu et al., 2016). Typically, the r-value of such correlations range between -0.3 and -0.4, meaning that the strength of the upper limb EVR predicts ∼10-15% of the variance of reach reaction time (Gu et al., 2016; Kozak & Corneil, 2021). We observed an even stronger negative correlation in the lower limb (average r-value of -0.58), but we note that this relationship is between the EVR magnitude on the *stance* leg and the RT of lift off of the *stepping* leg. Our force plate data in the anterolateral stepping condition shows that the likely kinetic consequence of stance-side GM recruitment and stepping-side GM suppression during the EVR is a rapid increase in ground reaction force generated by the stance leg and unloading of the stepping leg within less than 150 ms of target presentation, which propels the CoM towards the direction of the target. This is remarkably rapid and clearly preceded the APAs as generated in the medial stepping condition at ∼170 ms post stimulus presentation.

#### Muscle considerations

The current study underlines the importance of lateralized muscle activity in order to discern robust EVR detection through time-series ROC analysis. Target-selective muscle activity is a key feature of the EVR, as this provides evidence that the brain accounts for the target location in its visuomotor transformations, as opposed to a stereotyped response elicited, for example, by startling stimuli. While TA did show early muscle onsets (i.e. generally within the EVR window), it did so bilaterally, thus not meeting this criterion. EVRs in upper limb muscles are spatially tuned (Gu et al., 2019; Selen et al., in press), and it may be the case that TA activity will be lateralized for other target locations. Regardless, TA was unsuited to study lower limb EVRs in the current study context. In contrast, the experimental task required lateralized activity of GM for both the APA and the stepping movement, but in an opposite way and with different timing, which allowed us to study their interaction. As EVR prevalence in the lower limbs is now established, future studies should look into EVR expression in other lower-limb muscles. For example, it is plausible to assume that EVRs would also be present in muscles on the stepping side, where they could facilitate step execution (e.g. by expediting the lifting of the leg). A suitable candidate might be tensor fascia latae, which contributes to both hip flexion and abduction and previously demonstrated short-latency recruitment (98 ms) during on-line step adjustments in response to target shifts (Reynolds & Day, 2005).

### EVRs facilitate rapid stepping movements in the anterolateral stepping condition, but are detrimental in the anteromedial stepping condition

We demonstrated that the combination of stance width and target location, which required anterolateral or anteromedial steps had a major influence on the postural demands of the stepping task. As emphasized before, step initiation usually starts with an APA phase that ensures that balance demands are met prior to the subsequent stepping movement (Honeine et al., 2016). Previous work by Inaba and colleagues (2020) showed that APA size generally decreases as the stepping direction becomes more anterolateral (with stepping directions of up to 90° relative to a forward step). In comparison, the stepping direction in the anterolateral stepping condition in the current paradigm only deviated 11° from a forward step. Strikingly, we discovered that APAs were not only decreased, but generally not expressed in anterolateral stepping. APAs were strongly expressed in anteromedial stepping at a latency of ∼150ms after target appearance.

In conjunction with this finding, we found that the preceding EVR activity in GM corresponded to stepping-related activity and not APA-related activity. From a biomechanical point of view, this indicates that EVRs were favorable during anterolateral stepping and detrimental during anteromedial stepping. Because APAs apparently were not needed for maintaining balance in the anterolateral stepping condition, the observed EVR activity in stance side GM allowed for an extremely rapid step initiation, by directly propelling the CoM towards the stepping side.

In contrast, APAs were essential prior to step initiation in the anteromedial condition to account for the increased balance demands. Thus, initially GM needed to be activated on the ipsilateral side to shift the CoM towards the stance limb. The occasionally observed stepping-related EVR activity in stance-side GM *prior to the APA* was therefore detrimental to postural stability. Thus, following EVR expression in this high-postural demand condition, larger APAs were required to compensate for the EVRs. Due to the present study design, participants knew in advance whether an anterolateral or an anteromedial step needed to be performed. Higher-order cortical areas could therefore account for the increased balance demands in the anteromedial-target condition by a-priori downregulating the EVRs, as they would otherwise negatively affect step initiation. However, on trials where this cortical inhibition momentarily lapsed, the subsequently ensuing strong EVR hindered a fast stepping response towards the target, as postural demands needed to be met first.

These findings complement an earlier study from Nonnekes et al. (2010), which investigated on-line stepping adjustments in response to either lateral or medial target jumps in stroke patients. Results showed that the correction speed in medial mid-flight adjustments was significantly slower compared to lateral stepping adjustments, suggesting that the increased balance demands in the medial condition compelled the stroke patients to suppress medial stepping adjustments in order to safely complete the step. A similar comparison can be drawn to studies investigating the fast visuomotor network during either pro-reaches (towards the target) or anti-reaches (away from the target) (Gu et al., 2016; Kozak et al., 2020). In those studies, stronger EVRs preceded faster RTs in pro-reaches, but slower RTs in anti-reaches. Taken together, these results support the notion that the fast visuomotor system prioritizes the rapid goal-directed movement *towards* the target, while the experimental context (i.e. reaching away from the target during reaching or increasing postural demands during stepping) can dampen EVR expression. In situations where the dampening of EVR expression fails, subsequent phases of muscle recruitment have to be increased in order to correct for the kinetic consequences of the inappropriate EVR.

### Neural correlates of EVRs and postural control

Although the present paradigm taps into a subcortical fast-latency pathway evoking the EVR, the influence of higher-order cortical areas on the subcortical pathway via top-down processes should not be underestimated. In fact, specific features of the emerging target paradigm proposed by Kozak and colleagues (2020) engage higher-order processes, such as implied motion (Krekelberg et al., 2005) and temporal predictability (Contemori et al., 2021a) that help potentiate EVRs. Multiple studies have shown that top-down processes can significantly influence the properties of the EVRs. For example, EVR characteristics are influenced by expectations or cueing of target location, the immediacy of movement, hand-path barriers, or instructions in how to respond to the reaching target (Contemori et al., 2021a, 2021b, 2022a, 2022b; Gu et al., 2016, 2017; Kozak et al., 2020; Wood et al., 2015). These findings are consistent with the hypothesis that EVRs are regulated via the midbrain superior colliculus and its downstream connections with the brainstem reticular formation (Boehnke & Munoz, 2008; Contemori et al., 2022b; Corneil et al., 2004; Corneil & Munoz, 2014; Glover & Baker, 2019). Here we demonstrate that the expectation of postural instability can suppress EVRs in the lower limb. APAs are also thought to be regulated via a complex cortical-subcortical interplay involving primary sensorimotor cortex, supplementary motor area (Jacobs et al., 2009; Richard et al., 2017), basal ganglia and brainstem (Takakusaki, 2017). In addition, evidence suggests that the reticulospinal tract plays an important role in postural control and particularly the execution of APAs (Drew et al., 2004; Massion, 1992; Takakusaki et al., 2016). The signals are then projected to interneurons and motoneurons via the reticulospinal tract (Schepens & Drew, 2004; Takakusaki et al., 2016). Thus, it is plausible to assume that both postural control and EVRs innervate GM via the reticulospinal tract. However, APA onset latencies were longer compared to the hyper-direct subcortical EVR responses (see Results). Suppression of the EVR response in high postural-demand conditions was therefore crucial in order to maintain balance. This suppression potentially takes place at the level of the reticular formation in the brainstem. Future studies should explore this further.

### Limitations and future research

Participants knew in advance about the postural demands of the task. The decrease in EVR prevalence and magnitude in the anteromedial condition could therefore be caused by higher-order brain areas suppressing the EVR network on a block-by-block basis. Future research could further investigate this cortical-subcortical interaction by presenting anterolateral and anteromedial targets in an intermixed way rather than separated by block. If the decrease in EVR prevalence and magnitude in anteromedial conditions indeed depends on prior knowledge of the increased balance demands of the task, multiple scenarios are possible in an intermixed paradigm: on the one hand, participants may prioritize balance over speed in which case the EVRs will be minimized in both the anteromedial and anterolateral stepping conditions. On the other hand, the initial response may be to step as quickly as possible until cortical areas intervene to produce an APA when stepping anteromedially, in which case the EVRs will appear strong in both anteromedial and anterolateral stepping conditions, but as a consequence will negatively impact stepping RTs in the anteromedial condition.

Our study presented anteromedial or anterolateral targets at a retinal eccentricity of 2.8° or 9.9°, respectively, relative to the line of sight. We believe that this difference in retinal eccentricity did not cause the general absence of EVRs to anteromedial targets. Evidence from the upper limb suggests that the EVR is tuned to target position relative to the hand, not relative to the current line of sight (Contemori et al., 2022b; Gu et al., 2017). Indeed, robust EVRs on upper limb muscles (Gu et al 2018; Kearsley et al 2022) or other related profiles of muscle recruitment (Cross et al. 2019) can be initiated by parafoveal visual events, providing the eye and hand are not initially aligned. Nevertheless, future studies should adjust the paradigm such that targets are presented at the same retinal eccentricity, or incorporate a range of potential target locations for both posturally stable or unstable starting locations. Holding on to a handrail could also provide another means of manipulating postural demands during medial stepping.

## Conclusions

Our results suggest an intricate relationship between ultra-rapid, albeit posturally-dependent expression of EVRs, online control of the APA, and subsequent step initiation. The emerging target paradigm evoked strong and reliable EVRs in the hip abductor muscle gluteus medius on the stepping side in a low postural-demand condition, which facilitated rapid step initiation. Such strong EVRs coincided with an absence of APAs. However, as postural demands were increased, APAs became essential in order to maintain postural stability during step initiation. In this condition, stance-limb EVRs were largely absent. On trials where EVR suppression temporarily lapsed, EVRs disrupted postural stability, which necessitated larger APAs and longer stepping reaction times. We suggest that regulation of the putative subcortical EVR network is mediated by higher-order cortical areas. This top-down modulation is dependent on the expectation of postural (in)stability of the upcoming step. The successful adaption of the emerging target paradigm into a stepping task greatly increases the potential to further investigate this interaction between various neural networks in future studies across different disease states and across the lifespan.

## Additional Information

### Data availability statement

The authors confirm that the data supporting the findings of this study are available within the article and its supplementary materials.

### Author Contributions

**Lucas Billen:** Conceptualization, Methodology, Software, Formal analysis, Investigation, Data curation, Writing – Original Draft, Review & Editing, Visualization; **Brian Corneil:** Conceptualization, Methodology, Software, Resources, Writing - Review & Editing, Supervision, Funding acquisition; **Vivian Weerdesteyn:** Conceptualization, Methodology, Resources, Writing - Review & Editing, Supervision, Project administration, Funding acquisition

## Acknowledgements

We thank Drs. Tim Carroll, Gerald Loeb, and Guy Wallis, as well as members of the Corneil and Carroll lab, for comments on an earlier version of this manuscript. Thank you also to Maartje Meijer for helping in the early stages of the project.

## Supplementary material

### The influence of target appearance on EVR characteristics

Similar to the flashed target condition, EVR prevalence in the moving target condition was high for anterolateral stepping (16/16 participants) and low for anteromedial stepping (4/16 participants).

We investigated the effect of target appearance (flashed vs moving) on EVR latency. Since the anteromedial target condition did not yield discrimination times in every participant, we restricted the analysis to the anterolateral target condition. EVR latency was significantly longer in the flashed condition (*M* = 112 ms, *SD* = 6.83) compared to the moving condition (*M* = 108 ms, *SD* = 7.21; *t*(15) = 2.59, *p* = .020, Hedges’ *g* = 0.57).

A second t-test was performed to investigate the effect of target appearance on EVR magnitude. Here we also included the anteromedial condition. In the anterolateral conditions, moving targets resulted in significantly higher EVR magnitudes (*M* = .127, *SD* = 0.06) compared to flashed targets (*M* = .120, *SD* = 0.05; *t*(15) = 2.32, *p* < .035, Hedges’ *g* = 0.57). In the anteromedial conditions, there were no significant EVR magnitude differences between flashed and moving targets (*p* > .05).

Thus, our findings indicate that moving targets lead to faster and stronger EVRs compared to flashed targets. This contradicts the findings of Contemori et al. (2021), who reported lower EVR latency and higher EVR magnitude in reaches towards transient flashed targets compared to sustained moving targets. However, several methodological deviations may be responsible for these contradictory findings, such as the duration of the flashed target (8 ms in the Contemori study, versus 48 ms here) and the direction of the moving target (downwards in the Contemori study, sidewards here). The minor difference in EVR latency and EVR magnitude between stepping towards flashed targets and moving targets might also be the result of a slightly different stepping direction between the conditions. Since the moving target moved towards the edges of the treadmill, participants had to make more lateral steps towards moving targets compared to flashed targets.

Summary figures for all participants. For figure description, see **Figure 2**.

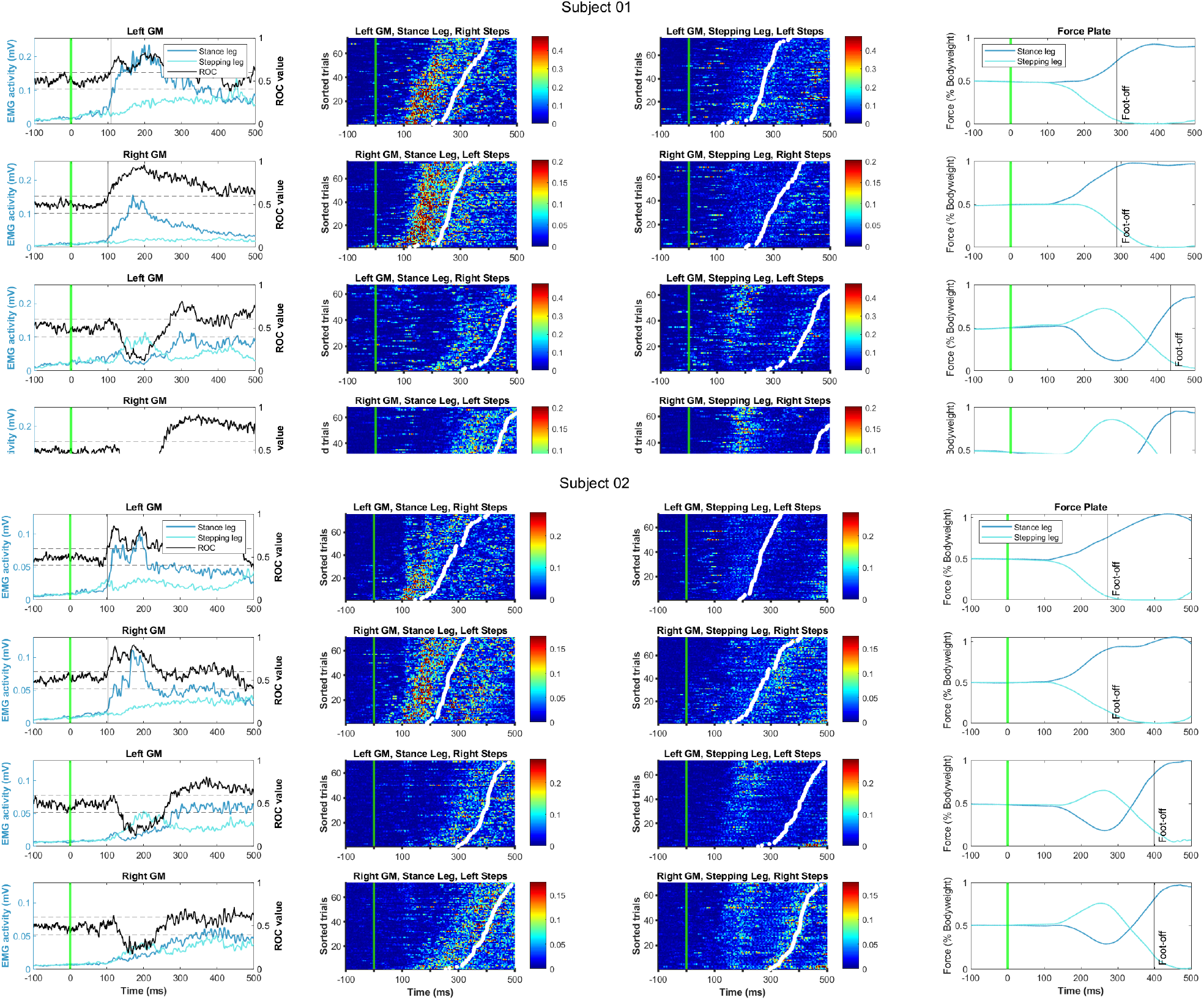
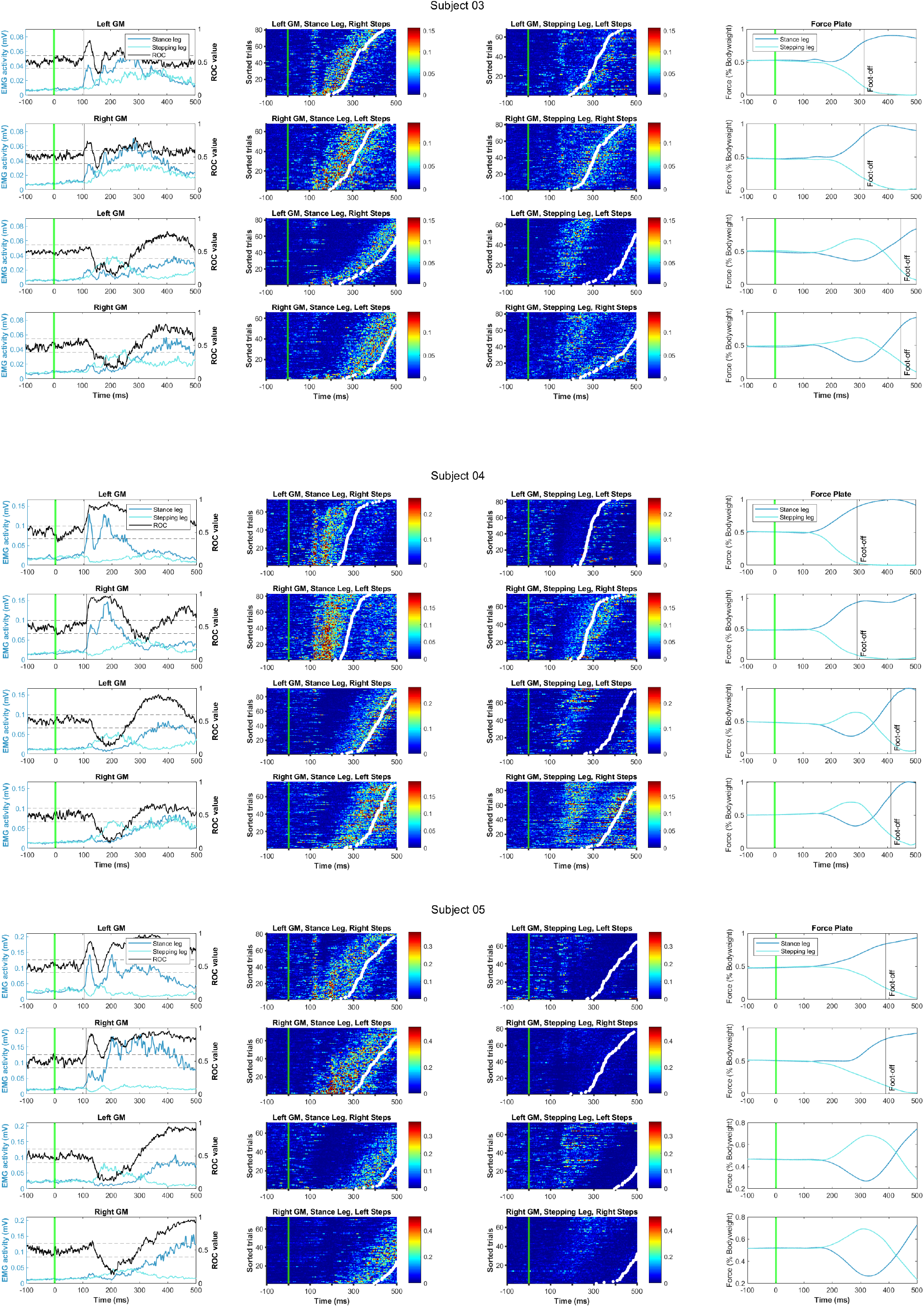
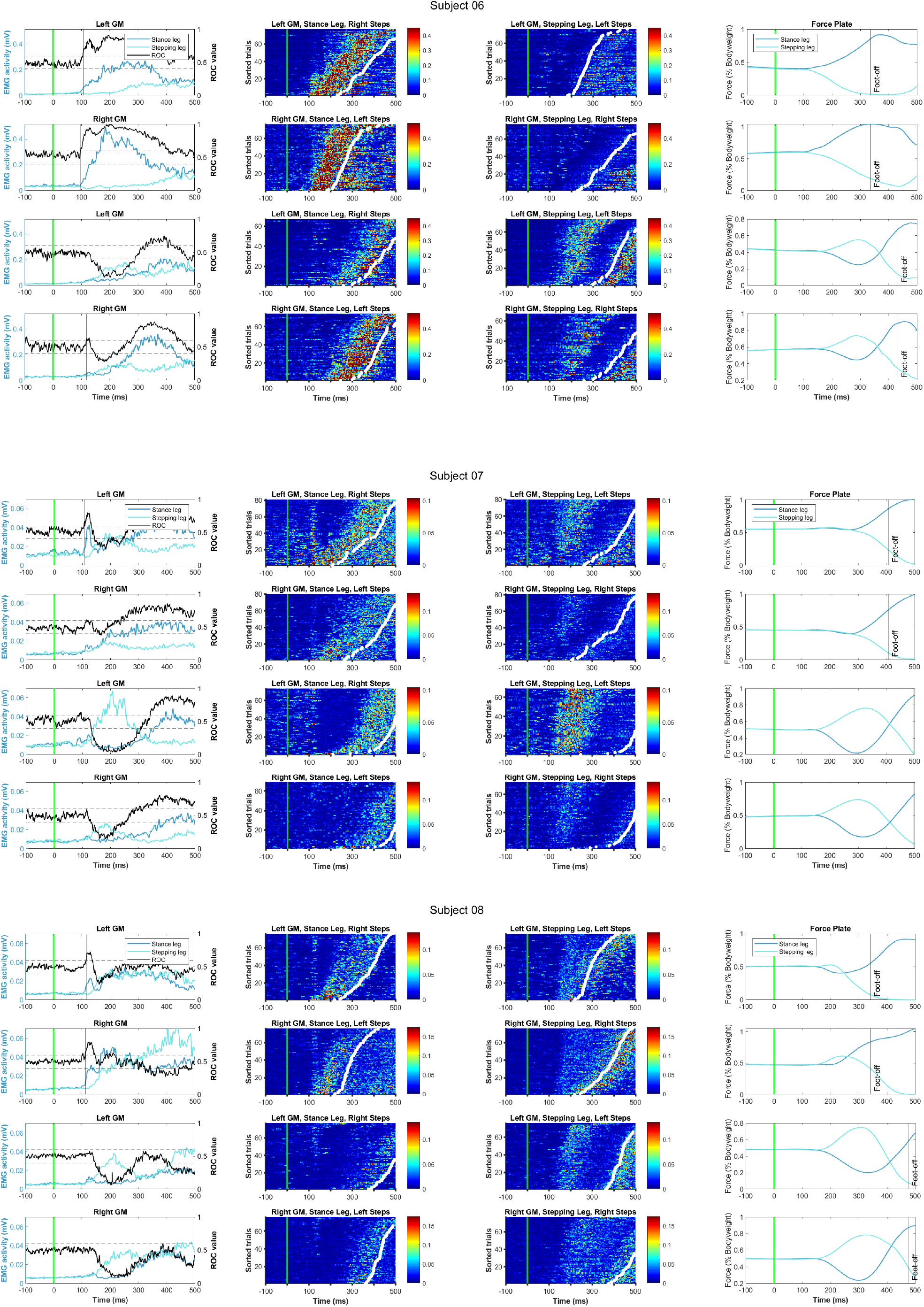
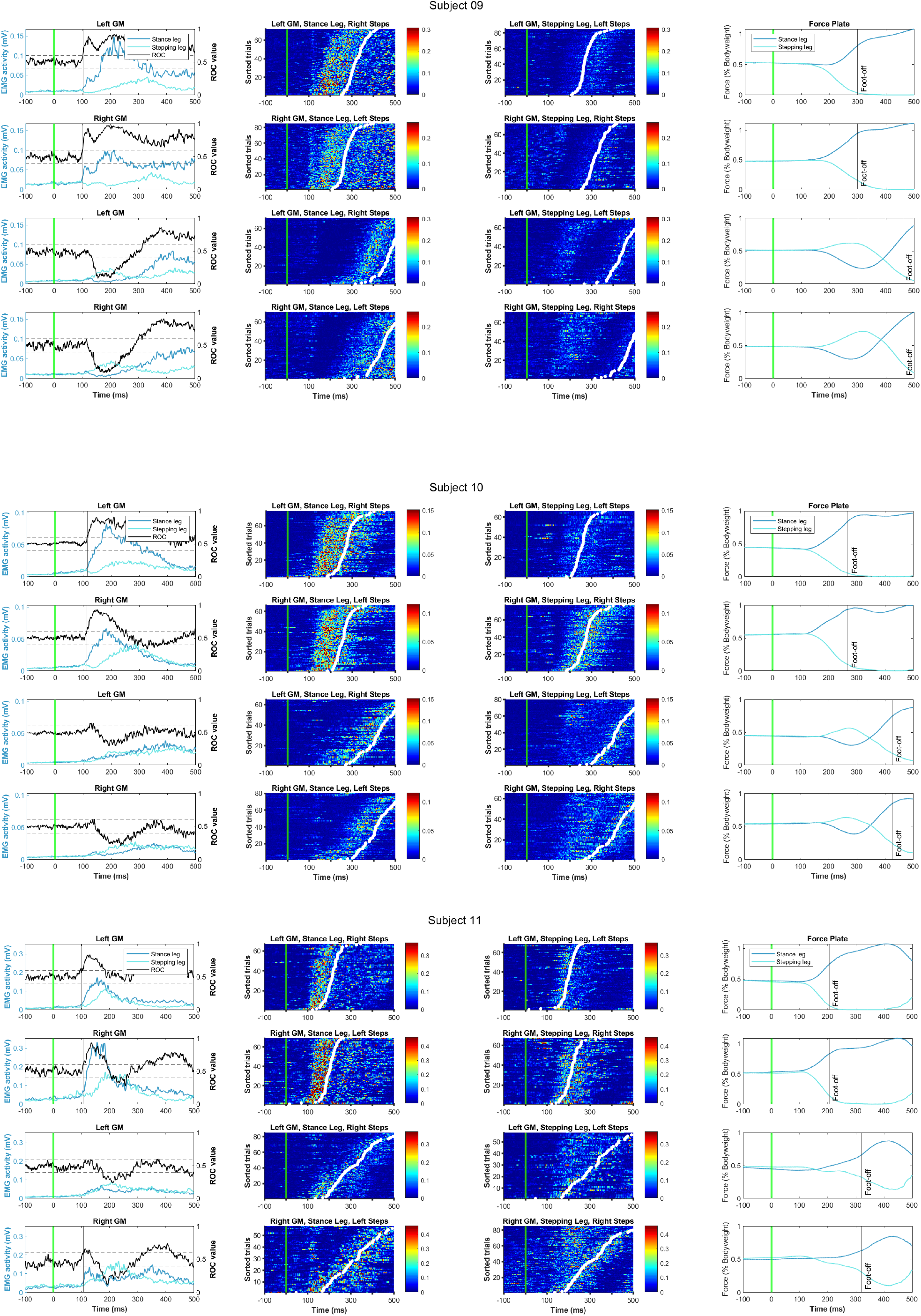
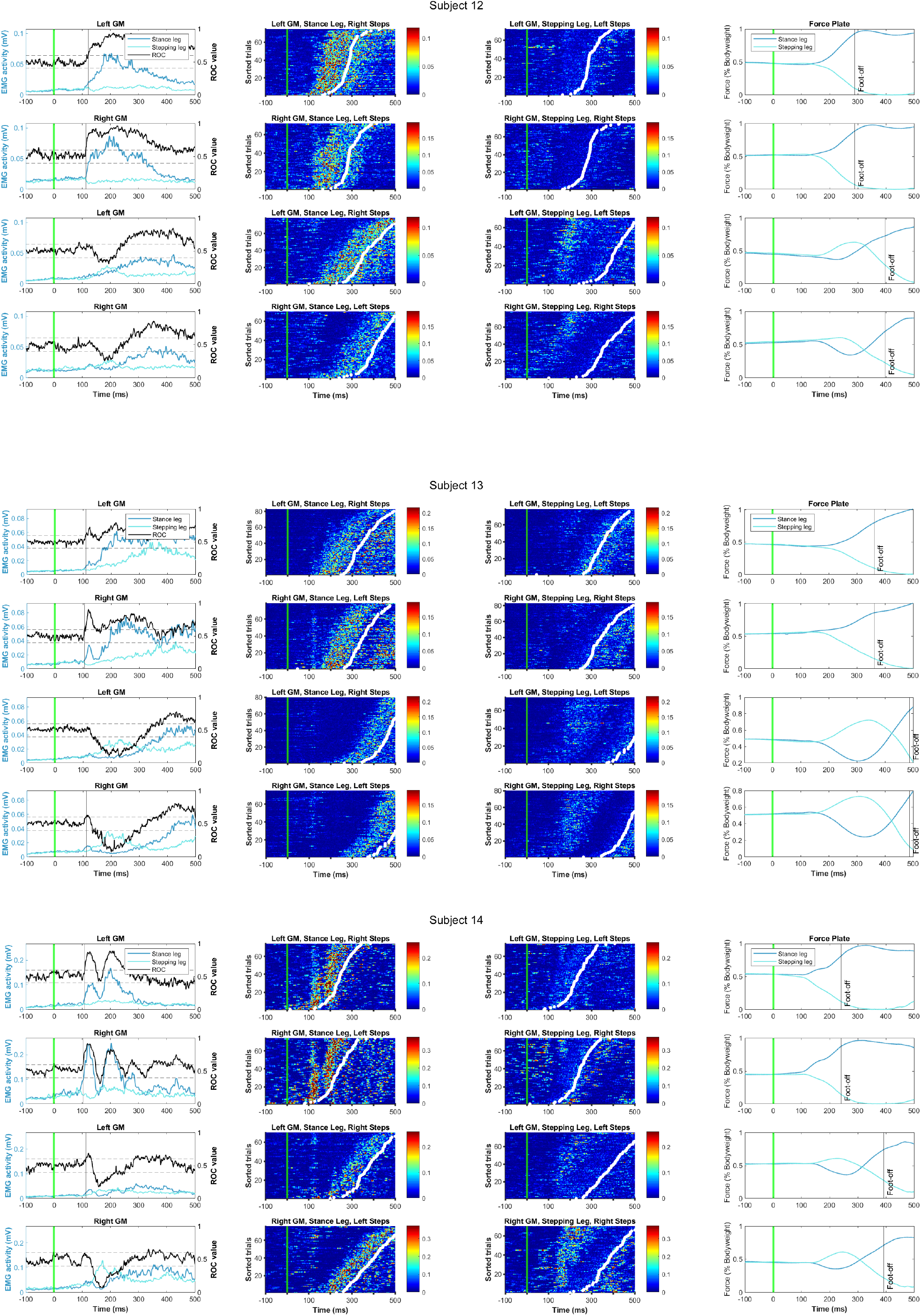
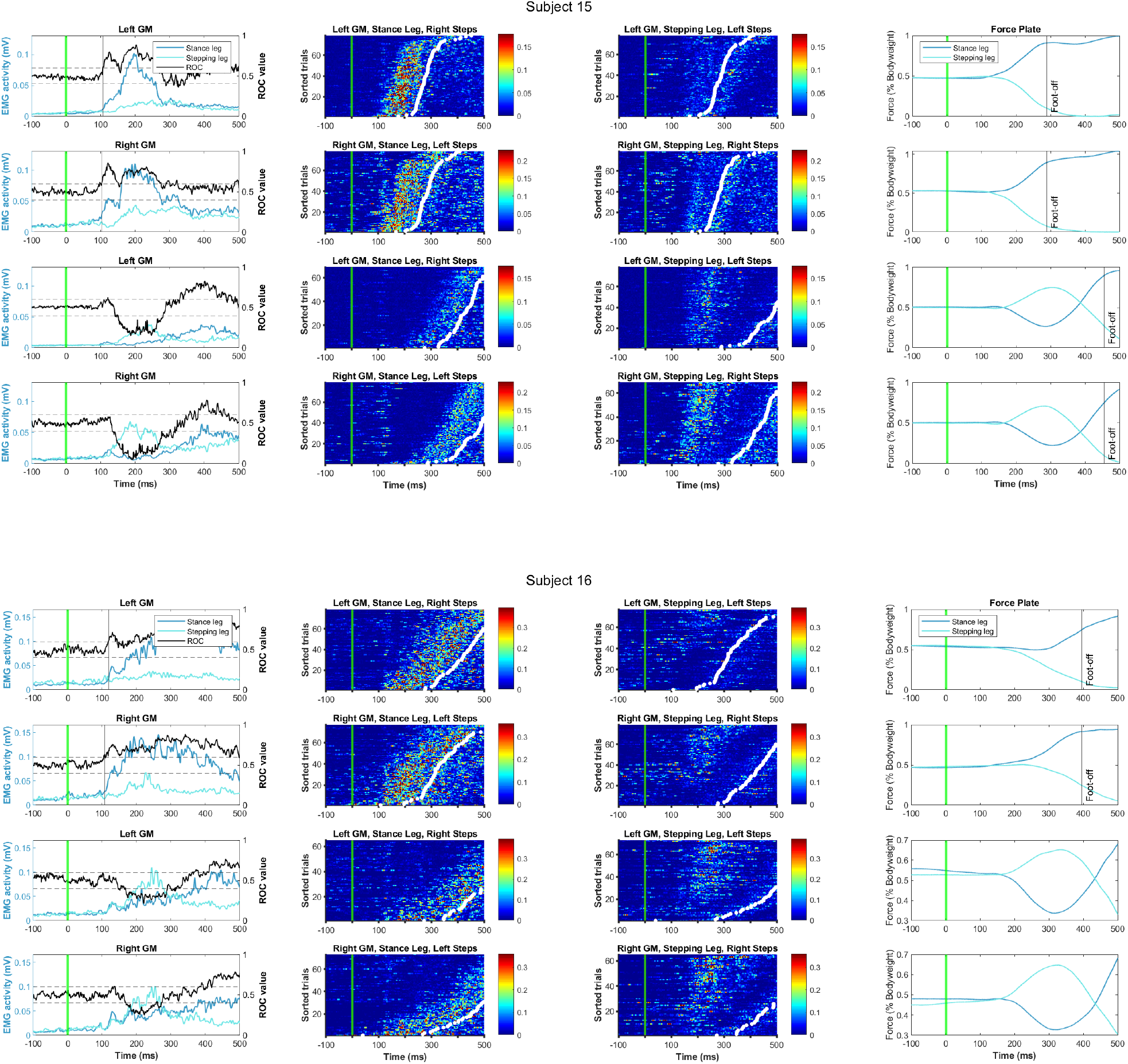

